# The impact of viral transport media on PCR assay results for the detection of nucleic acid from SARS-CoV-2 and other viruses

**DOI:** 10.1101/2020.06.09.142323

**Authors:** P.D. Kirkland, M.J. Frost

## Abstract

During the 2020 SARS-Cov-2 pandemic, there has been an acute shortage of viral transport medium. Many different products have been used to meet the demands of large-scale diagnostic and surveillance testing. The stability of SARS-Cov-2 RNA was assessed in several commercially produced transport media and an in-house solution. Coronavirus RNA was rapidly destroyed in the commercial transport media though the deleterious effects on intact virus were limited. Similar results were obtained for a Type A influenza virus. There was reduced detection of both virus and nucleic acid when a herpesvirus sample and purified DNA were tested. Collectively these data showed that the commercial viral transport media contained nucleases or similar substances and may seriously compromise diagnostic and epidemiological investigations.

Recommendations to include foetal bovine serum as a source of protein to enhance the stabilising properties of viral transport media are contraindicated. Almost all commercial batches of foetal bovine serum contain pestiviruses and at times other bovine viruses. In addition to the potential for there to be nucleases in the transport medium, the presence of these viruses and other extraneous nucleic acid in samples may compromise the interpretation of sequence data. The inclusion of foetal bovine serum presents a biosecurity risk for the movement of animal pathogens and renders these transport media unsuitable for animal disease diagnostic applications. While these transport media may be suitable for virus culture purposes, there could be misleading results if used for nucleic acid-based tests. Therefore, these products should be evaluated to ensure fitness for purpose.

## Introduction

For several decades a variety of medium solutions have been recommended to stabilise specimens for the detection of bacteria and viruses, particularly during diagnostic investigations. These have usually been based on balanced salt or saline solutions with a buffering capacity to maintain a “near-neutral” pH. To enhance the stability of viruses a spectrum of protein supplements has and continues to be recommended (1, 2, 3, 4). While some laboratories have prepared viral transport medium (VTM) “in house”, commercial preparations are used extensively and are often supplied as part of a sample collection kit with sterile swabs. Testing of samples by cultural methods meant that the emphasis of studies for the evaluation of these products originally focussed on the capacity of a preparation to maintain the infectivity of viruses at different temperatures while being held prior to and during transport and while being stored at the laboratory (5, 6). With the widespread introduction of molecular based diagnostic assays, especially real time PCR (qPCR), studies have been undertaken to evaluate the stability of viruses in VTMs, particularly in commercially prepared products, while being held at a range of temperatures (7, 8, 9). However, while thermal stability has been considered, generally little attention has been given to other characteristics of the VTM or the potential impact of endogenous components. One commercially available product is specifically designed to inactivate viruses and bacteria and contains components to inhibit the activity of nucleases that may be present in the sample (10, 11). There are some other products that are recommended for use in molecular detection assays but the manufacturers provide no comment that these products are unlikely to be suitable for samples where virus culture will be attempted.

During large-scale disease epidemics there can be pressure placed on the capacity of manufacturers to supply transport media and, during a pandemic, supply-chain and manufacturing pressures can become prohibitive. During the current (2020) SARS-Cov-2 pandemic, there has been an acute shortage of VTM in Australia because of a combination of both local and international demand, the lack of a local manufacturer and partly because of reduced international airline flights to Australia. Consequently, many different VTMs and similar solutions have been used to meet the demand for transport media generated by large scale diagnostic and surveillance testing. After becoming aware of concerns of variable results in some assays, we initiated a preliminary study to compare the stability of SARS-Cov-2 RNA in several commercially available VTMs and an in-house product. When disturbing results were found in a preliminary SARS-CoV-2 study, the investigation was extended to examine whether there was also an adverse impact on qPCR results for other viruses. Also of concern are recommendations (3, 4) to include foetal bovine serum (fbs) as a source of protein to enhance the stabilising properties of VTMs. This report documents observations of the adverse impact of certain VTMs on real time reverse transcription PCR (qRT-PCR) assays for the detection of SARS-CoV-2 virus as well as on a Type A influenza virus and a herpesvirus and discuss the broader implications of the inclusion of foetal bovine serum as a protein supplement to VTMs.

## Materials and Methods

### Samples

During the initial investigation, purified RNA from an Australian isolate (WMD DC1) of SARS-CoV-2 was supplied to the Elizabeth Macarthur Agriculture Institute (EMAI) by the Institute of Clinical Pathology and Medical Research (ICPMR), Westmead, New South Wales (NSW). A 1/1000 dilution of this RNA was made in tRNA (Sigma, 40ng/ul) to use as a reference preparation (designated R7215) and stored frozen at approximately −80°C. Subsequently, to include a sample which was expected to contain a high titre of intact, likely infectious virus, a sample from a recently diagnosed patient was selected (P66). This sample had also been stored frozen at approximately −80°C soon after it had been tested a few days earlier. Dilutions of these samples were tested in parallel (see below).

To extend the study beyond SARS-COV-2 virus, a Type A influenza virus, (L998, an isolate of H10N7 influenza (12)), was used. RNA was purified from L998 using standard methods (see below) and was diluted 1/1000 in the extraction elution buffer to prepare a working stock. Dilutions of this purified nucleic acid preparation were also tested in parallel with the infectious reference virus L998 as described for the SARS-CoV-2 virus.

Finally, a preparation of bovine herpesvirus 1 (BHV-1, M631) was similarly prepared as both purified DNA and infectious virus to determine the influence of transport media on a DNA virus.

### Viral transport media

The study included 4 different VTMs. VTM-1 was an “in-house” product based on phosphate buffered saline (PBS pH 7.2) supplemented with 0.5% gelatin (PBGS), antibiotics and 0.004% phenol red as a pH indicator. The other three were commercially manufactured VTMs – VTM-2 and VTM-3 are widely used, both in Australia as well as many countries in the northern hemisphere; VTM-4 is a new Australian manufactured product that is undergoing evaluation. VTM-2 and VTM-3 are believed to be supplemented with bovine serum albumen and gelatin and VTM-4 with foetal bovine serum. Sterile phosphate buffered saline (PBS), pH 7.2 was used as a control solution.

### Study design and sample/VTM mixture

For each virus under study, the concentrated reference viruses and the respective purified nucleic acids were diluted in either phosphate buffered saline (PBS) or one of the VTM solutions and were then subjected to nucleic acid extraction and testing by qPCR. Specifically, dilutions of the purified SARS-CoV-2 RNA (R7215) were prepared by adding either 50uL of R7215 to 450uL for a 1/10 dilution of the VTM under study or, for 1/2 dilutions, 100uL of diluted R7215 to an equal volume of the respective VTM. In the pilot experiment, only VTMs 1-3 were included. Within 15 minutes of preparation of the dilutions of RNA in each VTM, the diluted samples were extracted and tested by qRT-PCR as described below. Subsequently, this pilot experiment was repeated with the inclusion of PBS and VTM-4, as well as a series of dilutions of patient sample P66. These dilutions of RNA and virus were then extracted within 45 minutes of preparation and tested in several SARS-CoV-2 qRT-PCR assays. Later, less dilute preparations of R7215 were made so that the impact of the VTMs on higher concentrations of SARS-CoV-2 RNA could also be assessed. To obtain sufficient VTM as a diluent for this study, the contents (2.5-3mL) of 2 vials of VTM were pooled and mixed.

To investigate whether the results obtained from the experiments involving SARS-CoV-2 were applicable to other viruses, the same experimental design was applied to a Type A influenza virus and a herpesvirus and the respective purified viral nucleic acid extracts. The starting concentrations of nucleic acid and whole virus were adjusted so that the levels of nucleic acid (based on cycle-threshold (Ct) values) were very close to those used in the SARS-CoV-2 experiments. In each instance each diluted sample (nucleic acid or whole virus) was extracted and tested within 45 minutes of preparation of the dilutions.

As well as testing the whole virus & purified nucleic acid combinations, the impact of the different VTMs on the exogenous RNA internal control (XIPC) were also assessed by adding 1,000 or 10,000 copies of the XIPC to 100uL of each VTM and testing immediately or after the dilutions had been held at 25°C for 48 hours.

Finally, to confirm that VTM-1 can provide a high level of stability for clinical samples, 15 swabs from SARS-CoV-2 infected patients were removed from the transport medium in which they had been collected and were placed in new vials containing 1ml of VTM-1 and were then held at 4°C for 46-49 days before testing.

### Nucleic acid extraction and PCR assays

Total nucleic acid was extracted from 50uL of each sample (including dilutions of the already purified RNA or DNA) with a magnetic bead-based viral RNA extraction kit (MagMax96 Viral RNA – Ambion) that is run on a Kingfisher-96 magnetic particle handling system (ThermoFisher). After purification the nucleic acid was eluted in 50uL of kit elution buffer and 5uL run in the appropriate qRT-PCR. The primers and probe were directed at the SARS-CoV-2 E gene (13) and the IP4 assay targeting the RdRp gene (14) as developed by the Institut Pasteur (IP), Paris, France and recommended by the World Health Organisation. The SARS-CoV-2 primers and probe were used in a triplex assay format with the inclusion of the exogenous RNA internal control (XIPC) assay (15). The probes for the coronavirus assays (E/IP4) were both labelled with a FAM reporter and black hole quencher 1 (Biosearch Technologies) and the XIPC probe with a Vic reporter and TAMRA quencher (Life Technologies) as these combinations had been shown to provide high analytical sensitivity. A commercial reverse transcriptase mastermix (AgPath-ID One-Step RT-PCR kit -Life Technologies) was used and run, with slight modifications, under the standard cycling conditions recommended by the manufacturer, using an ABI7500 or Quantstudio 5 (Thermofisher Scientific) thermocycler. The combined annealing and extension temperatures were adjusted to 58°C and the assay was run for a total of 45 cycles. The baseline was manually set between 3 to 15 cycles and the threshold at 0.05. The results were expressed as cycle-threshold (Ct) values, being the number of PCR cycles at which the amplification plot crossed the threshold.

For comparative purposes a selection of samples was also tested using the CDC designed assay (2019-nCoV_N2) targeting the N gene, which was also run under the published conditions (16) with the same commercial mastermix.

Published primer and probe sets were used for the pan-reactive Type A influenza qRT-PCR (17), the BHV-1 qPCR (18) and the pan-pestvirus qRT-PCR (19) assays, each in a duplex format with the XIPC assay. Each assay utilised the AgPath-ID One-Step RT-PCR mastermix and were run for 45 cycles under the standard cycling conditions recommended by the manufacturer, with the baseline set automatically and the threshold at 0.05.

All assays included at least 2 positive controls, one negative control (tRNA) and a ‘no template’ control (NTC – nuclease free water). The XIPC RNA (approximately 80 copies/uL) was included in the sample lysis buffer prior to the extraction of nucleic acid from all test and control samples except for the NTC wells.

## Results

The results of these investigations are documented in Tables 1-9. The preliminary investigation (Table 1) showed that no SARS-CoV-2 RNA was detected after dilution inVTM-2 or VTM-3. In contrast, the results for VTM-1 were at the expected levels and those for the XIPC were highly reproducible for each sample, indicating that there had been efficient nucleic acid extraction and that there were no inhibitors of the qRT-PCR in the samples.

**Table 1.** Preliminary comparison of SARS-CoV-2 qRT-PCR Ct values when testing SARS-CoV-2 purified RNA diluted in different viral transport media.

| Sample | Dilution | VTM-1 |  | VTM-2 |  | VTM-3 |  |
| --- | --- | --- | --- | --- | --- | --- | --- |
|  |  | Target | XIPC | Target | XIPC | Target | XIPC |
| 1 | RNA R7215 @ $1/4 \times 10^{-3}$ | 30.48 | 29.60 | - | 29.51 | - | 29.36 |
| 2 | RNA R7215 @ $1/4 \times 10^{-4}$ | 34.56 | 29.74 | - | 29.52 | - | 29.64 |
| 3 | RNA R7215 @ $1/8 \times 10^{-4}$ | 39.04 | 29.34 | - | 29.48 | - | 29.24 |
| 4 | RNA R7215 @ $1/16 \times 10^{-4}$ | 41.89 | 29.97 | - | 29.44 | - | 29.5 |
| 5 | RNA R7215 @ $1/32 \times 10^{-4}$ | - | 29.32 | - | 29.82 | - | 29.43 |
| 6 | RNA R7215 @ $1/64 \times 10^{-4}$ | - | 29.60 | - | 30.07 | - | 29.22 |

\* - = not detected

To confirm the initial results, the experiment was repeated (Table 2, Samples 4-9), with the inclusion of both RNA and presumptively whole virus, as well as PBS and additional VTM solutions. VTM-1 and PBS gave very similar results for the RNA samples. Again, no SARS-Cov-2 RNA was detected in the dilutions prepared in VTMs 2-4. Consequently, a further set of dilutions (Table 2, Samples 1-3) were prepared in each solution to test whether there was any adverse effect on higher concentrations of RNA. SARS-Cov-2 RNA was not detected in any dilution prepared in VTM-2 and very weak reactivity (approaching the assays limit of detection) was observed for VTM-3 and VTM-4 in the sample with the highest concentration of RNA. In contrast, samples diluted in PBS and VTM-1 gave almost identical results, with a Ct value of approximately 21 for the highest RNA concentration. This difference between the PBS/VTM-1 result and the results for the other VTMs represents a reduction in analytical sensitivity of approximately 6 log_10_ for the detection of free SARS-CoV-2 RNA. The results for the XIPC were again highly reproducible and similar for each dilution in each solution, confirming the high efficiency of RNA extraction and no apparent impact of PCR inhibitors.

**Table 2.** Comparison of SARS-CoV-2 qRT-PCR Ct values when testing a broader range of dilutions of purified SARS -CoV-2 RNA in different viral transport media,

| Sample | Dilution | PBS |  | VTM-1 |  | VTM-2 |  | VTM-3 |  | VTM-4 |  |
| --- | --- | --- | --- | --- | --- | --- | --- | --- | --- | --- | --- |
|  |  | Target | XIPC | Target | XIPC | Target | XIPC | Target | XIPC | Target | XIPC |
| 1* | RNA R7215 @ $1 \times 10^{-1}$ | 20.9 | 28.4 | 20.8 | 28.2 | - | 30.1 | 39.4 | 31.1 | 39.8 | 30.6 |
| 2* | RNA R7215 @ $1 \times 10^{-2}$ | 24.3 | 29.7 | 24.3 | 29.6 | - | 30.5 | - | 30.9 | - | 30.4 |
| 3* | RNA R7215 @ $1 \times 10^{-3}$ | 25.8 | 30.5 | 25.6 | 30.0 | - | 30.5 | - | 30.3 | - | 30.7 |
| 4 | RNA R7215 @ $1/4 \times 10^{-3}$ | 29.4 | 30.3 | 30.3 | 30.2 | - | 30.3 | - | 29.9 | - | 28.8 |
| 5 | RNA R7215 @ $1/8 \times 10^{-3}$ | 31.8 | 29.5 | 31.3 | 28.3 | - | 29.5 | - | 30.7 | - | 30.2 |
| 6 | RNA R7215 @ $1/16 \times 10^{-3}$ | 32.5 | 29.8 | 31.6 | 28.7 | - | 30.0 | - | 30.5 | - | 29.3 |
| 7 | RNA R7215 @ $1/32 \times 10^{-3}$ | 33.8 | 29.9 | 34.6 | 29.0 | - | 30.8 | - | 30.5 | - | 30.9 |
| 8 | RNA R7215 @ $1/64 \times 10^{-3}$ | 40.2 | 29.9 | 40.3 | 28.4 | - | 30.6 | - | 29.0 | - | 29.1 |
| 9 | RNA R7215 @ $1/128 \times 10^{-3}$ | 40.3 | 27.9 | 40.8 | 30.0 | - | 29.0 | - | 29.6 | - | 30.4 |
| *Samples 1-3 tested 14 days after samples 4-9 |  |  |  |  |  |  |  |  |  |  |  |

Testing was then undertaken to determine whether there might be a reduction in sensitivity when testing a sample that presumptively contains high quality intact virions. In this instance, the results for the dilutions of virus (Table 3) were similar for each VTM and the PBS, as were the results for the XIPC. Similar results for both SARS-CoV-2 purified RNA and the patient sample were obtained in the CDC assay (2019-nCoV_N2) targeting the N gene (results not included).

**Table 3.** Comparison of SARS-CoV-2 qRT-PCR Ct values when testing a patient sample diluted in different viral transport media

| Sample | Dilution | PBS |  | VTM-1 |  | VTM-2 |  | VTM-3 |  | VTM-4 |  |
| --- | --- | --- | --- | --- | --- | --- | --- | --- | --- | --- | --- |
|  |  | Target | XIPC | Target | XIPC | Target | XIPC | Target | XIPC | Target | XIPC |
| 1 | P-66 @ $1 \times 10^{-2}$ | 23.7 | 29.7 | 24.1 | 29.9 | 24.2 | 30.0 | 25.0 | 30.3 | 24.5 | 29.5 |
| 2 | P-66 @ $1 \times 10^{-3}$ | 27.3 | 29.8 | 27.6 | 30.2 | 28.6 | 30.6 | 29.0 | 30.6 | 27.8 | 30.4 |
| 3 | P-66 @ $1 \times 10^{-4}$ | 28.0 | 29.6 | 29.9 | 31.0 | 32.2 | 30.1 | 32.7 | 31.2 | 31.0 | 30.4 |
| 4 | P-66 @ $1/2 \times 10^{-4}$ | 32.6 | 30.0 | 32.8 | 29.9 | 33.1 | 30.0 | 33.0 | 30.0 | 33.5 | 30.8 |
| 5 | P-66 @ $1/4 \times 10^{-4}$ | - | 30.4 | 37.8 | 30.0 | - | 30.5 | - | 30.6 | - | 29.6 |
| 6 | P-66 @ $1/8 \times 10^{-4}$ | - | 29.5 | - | 30.6 | - | 30.4 | - | 29.9 | - | 30.7 |
| 7 | P-66 @ $1/16 \times 10^{-4}$ | - | 30.8 | 40.1 | 30.2 | - | 30.6 | - | 30.2 | - | 31.0 |
| 8 | P-66 @ $1/32 \times 10^{-4}$ | - | 30.5 | - | 30.8 | - | 30.6 | - | 30.3 | - | 30.3 |
| 9 | P-66 @ $1/64 \times 10^{-4}$ | - | 29.6 | - | 30.5 | - | 29.9 | - | 31.0 | - | 30.3 |

Testing of the samples of VTM-1 to which swabs from SARS-CoV-2 positive patients had been added and stored at 4°C for 46-49 days gave results that were very similar to the original results. With the exception of a sample that had given a result (Ct=38.4) close to the limit of detection, the mean variation for the other 14 samples was very similar (0.6-1.0) Ct lower than the original result.

When the influenza virus and its homologous purified RNA were tested (Table 4), similar trends were noticed for the RNA results as had been observed for SARS-CoV-2. The results for VTM-1 and PBS were comparable but no RNA was detected in the dilutions in VTMs 2-4. The results for all solutions were similar at each dilution when whole virus was tested (data not included).

**Table 4.** Comparison of Type A Influenza qRT-PCR Ct values when testing purified influenza virus RNA diluted in different viral transport media

| Sample | Dilution | PBS |  | VTM-1 |  | VTM-2 |  | VTM-3 |  | VTM-4 |  |
| --- | --- | --- | --- | --- | --- | --- | --- | --- | --- | --- | --- |
|  |  | Target | XIPC | Target | XIPC | Target | XIPC | Target | XIPC | Target | XIPC |
| 1 | RNA L998 @ $1/2 \times 10^{-3}$ | 30.66 | 29.19 | 30.79 | 29.64 | - | 29.43 | - | 29.47 | - | 28.92 |
| 2 | RNA L998 @ $1/4 \times 10^{-3}$ | 31.71 | 29.49 | 31.33 | 29.50 | - | 29.11 | - | 29.09 | - | 29.59 |
| 3 | RNA L998 @ $1/8 \times 10^{-3}$ | 33.96 | 29.88 | 33.60 | 30.10 | - | 29.78 | - | 29.71 | - | 29.59 |
| 4 | RNA L998 @ $1/16 \times 10^{-3}$ | 34.52 | 29.57 | 34.02 | 29.48 | - | 29.65 | - | 29.69 | - | 29.23 |
| 5 | RNA L998 @ $1/32 \times 10^{-3}$ | 34.72 | 29.54 | 34.40 | 29.61 | - | 29.58 | - | 29.61 | - | 29.47 |
| 6 | RNA L998 @ $1/64 \times 10^{-3}$ | - | 29.28 | 37.49 | 29.49 | - | 29.51 | - | 29.23 | - | 29.35 |

Finally, to establish whether these effects were limited to testing of RNA viruses, a sample of bovine herpesvirus-1 was tested in the same model and range of dilutions. The results for each of the VTMs and PBS containing the highest concentrations of herpesvirus DNA were similar (Table 5) although there was a distinct trend towards higher Ct values for VTM-2 and VTM-3. The preparations of VTM-2 and VTM-3 with the lowest concentrations of DNA gave negative results. To confirm that the differences observed were not due to random variation around the limit of detection of the assay, eight replicates for the two lowest DNA concentrations (highest two dilutions of DNA) were then tested (Table 6). These results confirmed that there had been some impact of VTM-2, VTM-3 and VTM-4 on the stability of the purified DNA extracts. As had been noted for previous experiments, the XIPC gave highly reproducible results for all samples tested.

**Table 5.** Comparison of Bovine Herpesvirus −1 qPCR Ct values when testing purified BHV-1 DNA diluted in different viral transport media.

| Sample | Dilution | PBS |  | VTM-1 |  | VTM-2 |  | VTM-3 |  | VTM-4 |  |
| --- | --- | --- | --- | --- | --- | --- | --- | --- | --- | --- | --- |
|  |  | Target | XIPC | Target | XIPC | Target | XIPC | Target | XIPC | Target | XIPC |
| 1 | DNA M631 @ $1/2 \times 10^{-2}$ | 30.3 | 29.4 | 30.2 | 28.9 | 32.2 | 28.8 | 32.8 | 29.6 | 32.3 | 29.8 |
| 2 | DNA M631 @ $1/4 \times 10^{-2}$ | 31.5 | 29.5 | 31.6 | 29.4 | 33.4 | 29.1 | 32.8 | 29.0 | 33.5 | 29.5 |
| 3 | DNA M631 @ $1/8 \times 10^{-2}$ | 32.4 | 29.5 | 32.0 | 29.3 | 33.7 | 29.2 | 34.4 | 30.2 | 34.8 | 30.0 |
| 4 | DNA M631 @ $1/16 \times 10^{-2}$ | 32.9 | 28.9 | 33.7 | 29.0 | 36.8 | 29.7 | 34.8 | 29.6 | 34.7 | 29.9 |
| 5 | DNA M631 @ $1/32 \times 10^{-2}$ | 35.5 | 29.3 | 34.0 | 29.3 | 36.5 | 29.6 | 36.3 | 29.7 | 34.6 | 29.7 |
| 6 | DNA M631 @ $1/64 \times 10^{-2}$ | 36.3 | 29.3 | 34.8 | 29.5 | - | 29.7 | - | 29.5 | 35.9 | 29.2 |

**Table 6.** Bovine Herpesvirus −1 qPCR Ct values when testing replicates of purified BHV-1 DNA diluted in different viral transport media near the presumed limit of detection

| Sample | Dilution | PBS |  | VTM-1 |  | VTM-2 |  | VTM-3 |  | VTM-4 |  |
| --- | --- | --- | --- | --- | --- | --- | --- | --- | --- | --- | --- |
|  |  | Target | XIPC | Target | XIPC | Target | XIPC | Target | XIPC | Target | XIPC |
| 5 | DNA M631 @ 1/32 x 10<sup>-2</sup> | 34.2 | 29.6 | 35.6 | 29.8 | 37.0 | 29.6 | 36.7 | 29.8 | 36.5 | 29.3 |
| 5 | DNA M631 @ 1/32 x 10<sup>-2</sup> | 33.5 | 29.8 | 33.1 | 29.1 | 40.0 | 29.6 | - | 29.4 | 34.8 | 29.1 |
| 5 | DNA M631 @ 1/32 x 10<sup>-2</sup> | 34.4 | 29.6 | 34.6 | 29.1 | 37.8 | 29.3 | 35.4 | 29.4 | 36.7 | 29.6 |
| 5 | DNA M631 @ 1/32 x 10<sup>-2</sup> | 34.2 | 29.4 | 34.2 | 29.2 | 38.6 | 29.6 | 37.1 | 29.4 | 34.6 | 29.5 |
| 5 | DNA M631 @ 1/32 x 10<sup>-2</sup> | 34.3 | 29.8 | 34.0 | 29.3 | 39.9 | 28.7 | 37.5 | 29.7 | 35.4 | 29.0 |
| 5 | DNA M631 @ 1/32 x 10<sup>-2</sup> | 34.1 | 29.3 | 34.8 | 28.8 | 37.7 | 29.4 | 36.8 | 29.5 | 35.5 | 29.2 |
| 5 | DNA M631 @ 1/32 x 10<sup>-2</sup> | 33.7 | 29.1 | 33.9 | 28.9 | 36.5 | 29.5 | - | 28.9 | 35.2 | 29.7 |
| 6 | DNA M631 @ 1/64 x 10<sup>-2</sup> | 34.6 | 29.9 | 34.8 | 29.4 | - | 29.4 | 36.7 | 29.2 | 38.2 | 29.2 |
| 6 | DNA M631 @ 1/64 x 10<sup>-2</sup> | 34.3 | 29.2 | 36.0 | 28.9 | 40.8 | 29.4 | - | 29.5 | 35.9 | 29.6 |
| 6 | DNA M631 @ 1/64 x 10<sup>-2</sup> | 34.4 | 29.4 | 34.6 | 29.4 | 41.9 | 29.3 | - | 28.4 | 36.4 | 29.3 |
| 6 | DNA M631 @ 1/64 x 10<sup>-2</sup> | 33.6 | 29.6 | 36.1 | 29.4 | 43.9 | 29.6 | 36.5 | 29.8 | 37.7 | 29.2 |
| 6 | DNA M631 @ 1/64 x 10<sup>-2</sup> | 34.6 | 29.5 | 35.5 | 29.5 | - | 29.4 | - | 29.8 | 35.7 | 29.9 |
| 6 | DNA M631 @ 1/64 x 10<sup>-2</sup> | 35.3 | 29.5 | 35.0 | 29.5 | - | 29.6 | - | 29.4 | 37.5 | 29.0 |
| 6 | DNA M631 @ 1/64 x 10<sup>-2</sup> | 34.5 | 29.0 | 35.3 | 29.9 | - | 29.7 | - | 29.8 | - | 29.3 |

When the dilutions of bovine herpesvirus were tested, the adverse effects that had been observed with the purified DNA were again noted but perhaps in a more profound manner Table 7). Reduced sensitivity was observed with samples diluted in VTM-2, VTM-3 and VTM-4.

**Table 7.**
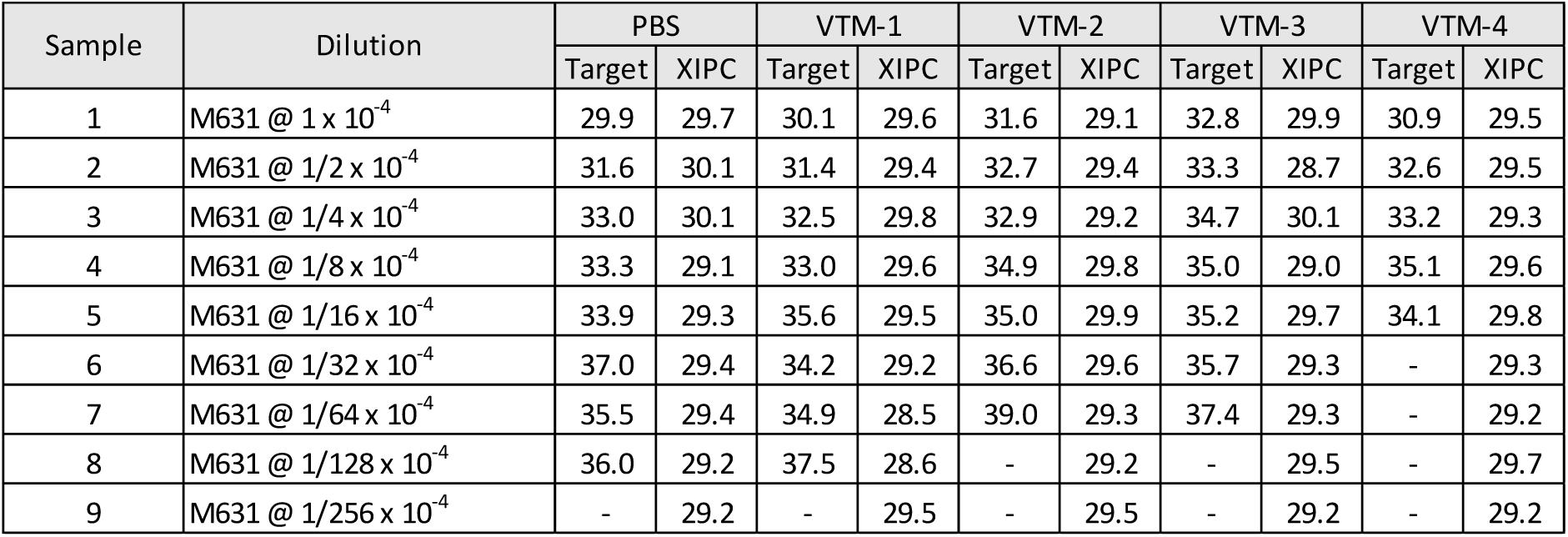
Comparison of Bovine Herpesvirus −1 qPCR Ct values when testing BHV-1 virus diluted in different viral transport media

| Sample | Dilution | PBS |  | VTM-1 |  | VTM-2 |  | VTM-3 |  | VTM-4 |  |
| --- | --- | --- | --- | --- | --- | --- | --- | --- | --- | --- | --- |
|  |  | Target | XIPC | Target | XIPC | Target | XIPC | Target | XIPC | Target | XIPC |
| 1 | M631 @ 1 x 10<sup>-4</sup> | 29.9 | 29.7 | 30.1 | 29.6 | 31.6 | 29.1 | 32.8 | 29.9 | 30.9 | 29.5 |
| 2 | M631 @ 1/2 x 10<sup>-4</sup> | 31.6 | 30.1 | 31.4 | 29.4 | 32.7 | 29.4 | 33.3 | 28.7 | 32.6 | 29.5 |
| 3 | M631 @ 1/4 x 10<sup>-4</sup> | 33.0 | 30.1 | 32.5 | 29.8 | 32.9 | 29.2 | 34.7 | 30.1 | 33.2 | 29.3 |
| 4 | M631 @ 1/8 x 10<sup>-4</sup> | 33.3 | 29.1 | 33.0 | 29.6 | 34.9 | 29.8 | 35.0 | 29.0 | 35.1 | 29.6 |
| 5 | M631 @ 1/16 x 10<sup>-4</sup> | 33.9 | 29.3 | 35.6 | 29.5 | 35.0 | 29.9 | 35.2 | 29.7 | 34.1 | 29.8 |
| 6 | M631 @ 1/32 x 10<sup>-4</sup> | 37.0 | 29.4 | 34.2 | 29.2 | 36.6 | 29.6 | 35.7 | 29.3 | - | 29.3 |
| 7 | M631 @ 1/64 x 10<sup>-4</sup> | 35.5 | 29.4 | 34.9 | 28.5 | 39.0 | 29.3 | 37.4 | 29.3 | - | 29.2 |
| 8 | M631 @ 1/128 x 10<sup>-4</sup> | 36.0 | 29.2 | 37.5 | 28.6 | - | 29.2 | - | 29.5 | - | 29.7 |
| 9 | M631 @ 1/256 x 10<sup>-4</sup> | - | 29.2 | - | 29.5 | - | 29.5 | - | 29.2 | - | 29.2 |

In order to further confirm that the differences observed between the different dilutions of virus were not due to random variation around the limit of detection of the assay, seven replicates of each sample from the last 2 dilutions that were positive in PBS were tested again. The results (Table 8) for both dilutions clearly confirmed the inferior performance of VTM-2 and VTM-3 with most replicates giving negative results or extremely high Ct values. VTM-4 gave acceptable results.

**Table 8.** Bovine Herpesvirus −1 qPCR Ct values when testing replicates of BHV-1 virus diluted in different viral transport media near the limit of detection.

| Sample | Dilution | PBS |  | VTM-1 |  | VTM-2 |  | VTM-3 |  | VTM-4 |  |
| --- | --- | --- | --- | --- | --- | --- | --- | --- | --- | --- | --- |
|  |  | Target | XIPC | Target | XIPC | Target | XIPC | Target | XIPC | Target | XIPC |
| 7 | M631 @ 1/64 x 10<sup>-4</sup> | 35.3 | 29.8 | 34.1 | 29.4 | 37.2 | 29.5 | 45.0 | 29.6 | 35.2 | 29.8 |
| 7 | M631 @ 1/64 x 10<sup>-4</sup> | 34.5 | 30.3 | 34.9 | 30.0 | 38.8 | 29.4 | 35.3 | 29.8 | 35.0 | 29.4 |
| 7 | M631 @ 1/64 x 10<sup>-4</sup> | 35.2 | 29.7 | 33.7 | 29.2 | 36.6 | 29.8 | - | 30.3 | 36.8 | 29.9 |
| 7 | M631 @ 1/64 x 10<sup>-4</sup> | 34.7 | 29.6 | 34.7 | 29.8 | 42.7 | 29.2 | 34.8 | 30.0 | - | 30.1 |
| 7 | M631 @ 1/64 x 10<sup>-4</sup> | 36.8 | 29.8 | 34.0 | 29.8 | 40.4 | 29.8 | - | 29.8 | 35.7 | 29.8 |
| 7 | M631 @ 1/64 x 10<sup>-4</sup> | 34.5 | 29.6 | 35.9 | 29.7 | 38.3 | 29.5 | 36.0 | 29.9 | NT | NT |
| 8 | M631 @ 1/128 x 10<sup>-4</sup> | - | 29.8 | 35.7 | 29.5 | 36.1 | 29.6 | - | 30.3 | 36.7 | 29.9 |
| 8 | M631 @ 1/128 x 10<sup>-4</sup> | - | 29.2 | 35.0 | 29.7 | - | 30.0 | - | 29.9 | - | 30.3 |
| 8 | M631 @ 1/128 x 10<sup>-4</sup> | 35.5 | 29.5 | 36.8 | 29.1 | - | 29.9 | - | 29.9 | 36.8 | 30.3 |
| 8 | M631 @ 1/128 x 10<sup>-4</sup> | 44.5 | 30.3 | - | 29.9 | - | 30.0 | NT | NT | NT | NT |
| 8 | M631 @ 1/128 x 10<sup>-4</sup> | 34.2 | 30.0 | 37.2 | 29.8 | - | 30.4 | - | 29.9 | 36.2 | 29.8 |
| 8 | M631 @ 1/128 x 10<sup>-4</sup> | 36.9 | 30.5 | 34.8 | 29.6 | - | 29.9 | 37.1 | 30.5 | 36.6 | 30.7 |

NT= not tested

The experiments undertaken have shown that some of the VTM solutions examined had a significant and deleterious impact on purified viral RNA and variable effects on DNA. The synthetic XIPC RNA that was used throughout this study was not affected because it was prepared in a tRNA solution and included in the sample lysis buffer which includes inhibitors of nuclease activity and the sample is extracted immediately after addition of the buffer. However, to confirm that the XIPC construct could be affected by components in the VTMs under study, two concentrations of XIPC were prepared in PBS and each of the VTM solutions and tested after being held at room temperature for 1 or 48 hours. The adverse effect of each commercial VTM (Table 9) was apparent within the first hour and no RNA was detected after 48 hours.

**Table 9.** qPCR-PCR Ct values for the exogenous synthetic RNA when diluted in different viral transport media and tested immediately and after holding at room temperature for 2 days

| Dilution | PBS | VTM-1 | VTM-2 | VTM-3 | VTM-4 |
| --- | --- | --- | --- | --- | --- |
| XIPC 10,000 copies, 1 hour at 25°C | 28.7 | 29.1 | 33.9 | 35.7 | 30.7 |
| XIPC 10,000 copies, 48 hours at 25°C | 28.9 | 29.2 | - | - | - |
| XIPC 1,000 copies, 1 hour at 25°C | 33.0 | 32.6 | 36.6 | - | 34.8 |
| XIPC 1,000 copies, 48 hours at 25°C | 32.3 | 32.5 | - | - | - |

## Discussion

The results of these studies clearly indicate that the commercially prepared VTM solutions have had an adverse impact on the ability to detect both SARS-CoV-2 and influenza RNA. The results for the RNA preparations diluted in the commercial VTMs would suggest that there are components of these VTMs that have prevented the detection of the RNA in these samples. As the detection of the XIPC was not affected, we propose that these results provide unequivocal evidence of the presence of a nuclease(s) or similar substance in these VTMs. The impact on these samples was rapid as all samples were extracted within one hour of preparation yet no RNA was detected in a sample that was estimated to contain more than 3000 copies of viral RNA in a 5uL sample. The same result was obtained with assays that were directed at 3 different regions of the SARS-CoV-2 genome. The results obtained for the virus samples that were diluted to similar concentrations as the RNA samples and held for the same time period also support this hypothesis because it is expected that these samples contained a high proportion of intact nucleocapsids that offered protection to the viral RNA. The same trends were observed with the Influenza A virus RNA and presumptively intact virions. Although the effects on the herpesvirus samples were less pronounced, nevertheless there was some impact on both purified DNA and whole virus.

It is also important to recognise that these observations reflect the outcome of contact between viral nucleic acid and VTM for less than 1 hour in each instance. The levels of nucleic acid that were destroyed after almost immediate addition to the VTMs were not insignificant. With a concentration of RNA that gave a Ct value of approximately 21 in both PBS and VTM-1, this represents a reduction of approximately ten thousand-fold and cannot be ignored. While it might be argued that the adverse impact on whole virus appeared to be slight, free nucleic acid and perhaps whole virus was destroyed at levels that could be of diagnostic relevance (20). While the impact on RNA viral genomes is likely to be markedly greater than on DNA sequences, the outcome cannot be predicted as secondary structure may also have an influence (20) and the speed and severity of the impact may vary depending on the nucleic acid target, as shown by the differences between the results for the two RNA viruses and the XIPC. Further, it cannot be assumed that the target nucleic acid will always be protected by nucleoprotein. Degradation will occur during the course of an infection and also under conditions where sample collection, transport and storage are sub-optimal. Additionally, the adverse effects observed in this study could potentially be exacerbated with alternative nucleic acid purification technologies that take longer than the 20 minutes required for this magnetic-bead based method. While undertaking surveillance and epidemiological tracing during a pandemic, failure to detect a moderate level of RNA in a person who is asymptomatic could result in a critical source of infection remaining undetected. However, with the selection of an appropriate transport medium, sample degradation, even at room temperature, can be minimal. This is clearly shown by the performance of the ‘in house’ medium (VTM-1) where there is little evidence of deterioration of the RNA samples after holding at room temperature for 2 days and no significant deterioration of virus when held at 4°C for more than 6 weeks.

The World Health Organisation, the US Centers for Disease Control and Prevention (CDC) and the UK government have each provided recommendations for the formulation of VTM solutions to be used for the collection of specimens for SARS-CoV-2 testing and comment that a supplement of protein or glycerol should be added to enhance the stability of viruses (2, 3, 4). We believe that, while this is an essential feature of a high quality VTM, it is in achieving this requirement that the current problem with the commercially available VTMs may have arisen. Our ‘in house’ VTM includes gelatin which is extracted from animal tissues by treatment at very low or high pH, and prolonged boiling at high temperatures before sterilisation and drying. These steps inactivate both enzymes and infectious agents that are present as well as destroying residual nucleic acid. Further, during the preparation of VTM-1, the PBS solution to which the gelatin has already been added is also sterilised by heat treatment. In contrast, products that include bovine serum albumen or other serum-derived components, such as VTMs 2, 3 and 4, cannot be sterilised by autoclaving without coagulation of the protein supplement. Therefore, for these VTMs, the raw materials must each be free of nucleases and proteinases prior to sterilisation by methods that do not include heating.

The choice of protein supplement is also an important consideration from the perspective of inadvertent addition of adventitious agents to the VTM and perhaps genomic DNA from animals from which the protein supplement has been derived. Foetal bovine serum (fbs) is a recommended supplement (3, 4) but almost all commercial batches of fbs contain pestiviruses and at times other bovine viruses. A moderate concentration of BVDV viral RNA (Ct=30.6, data not included) was detected in VTM-4. As whole genome nucleic acid sequencing is now often undertaken on many original, uncultured patient samples, the presence of these viruses and other extraneous nucleic acid in samples may reduce the sensitivity of sequencing protocols and complicate the interpretation of sequence data. The nucleases present may also have an impact on the quantity of RNA available for sequencing. Furthermore, the inclusion of fbs presents a biosecurity risk for the movement of animal pathogens between countries and renders such VTM solutions unsuitable for many animal disease diagnostic or research applications.

As indicated in the UK government guideline (4) there is a clear need for the “*Use (of) alternative swabs and transport medium in accordance with a locally validated laboratory strategy*” to demonstrate fitness for purpose. The specifications for products that might be used for both nucleic acid detection methods and virus culture are likely to be more rigorous than for those VTMs that are only used for one laboratory method. The special requirements for products that are suitable for nucleic acid testing have been recognised by some manufacturers who have developed specific transport media to inactivate the viruses of interest and to minimise the degradation of nucleic acid (10, 11). Some of the major manufacturers of VTM solutions also offer products with additives to reduce nuclease activity but most of these also preclude opportunities to undertake virus culture. However, these limitations are often not apparent to purchasing departments, especially during a pandemic, when any VTM may be mistakenly thought to be “fit for purpose”.

In conclusion, the results of this study provide examples of how the composition of a VTM could have an impact on the outcome of nucleic acid based testing and, in particular, situations where either there is a need to detect RNA that is not packaged into a nucleocapsid or where RNA constructs may be diluted in a VTM for use as a positive control in an assay or perhaps for proficiency testing. Finally, and particularly in the face of a pandemic, users should be reminded that products fit for one purpose may not be suitable for an alternative use. A product that may be eminently suitable for virus culture purposes could result in misleading results if used for nucleic acid-based tests.

## Acknowledgements

The authors are indebted to Mr Ian Carter, Institute of Clinical Pathology and Medical Research, Westmead Hospital, Westmead NSW for the generous supply of the purified SARS-CoV-2 RNA and for referral of the patient sample. We also appreciate the productive discussions with Drs Catherine Pitman and Dominic Dwyer, NSW Health Pathology regarding the need for rigorous evaluation of viral transport media. We also thank Dr Deb Finlaison for helpful comments on the draft manuscript and Shannon Mollica, Rodney Davis and other staff of the Virology Laboratory at EMAI for their assistance during the preparation of samples for the initial experiment and the longer-term storage of swabs.

